# Computational modelling of local fluctuations causing transient solvent exposure of protein amides

**DOI:** 10.1101/2022.05.19.492675

**Authors:** Soumendranath Bhakat, Pär Söderhjelm

## Abstract

Hydrogen exchange (HX) between protein amides and solvent water molecules can function as a probe for protein dynamics and provide a bridge between the experimental and computational worlds. However, it is important that the underlying assumptions are tested on well-known systems. Here we perform an analysis of a long MD simulation of BPTI, which has previously been used by *Persson and Halle* to propose a functional definition of the exchange-competent configurations. We find support for the hypothesis that the exchange-competent configurations typically do not constitute a metastable state *per se*, but rather occur as the outermost tail of the water distribution around a metastable *broken state* directly identifiable as having a broken intermolecular hydrogen bond and increased solvent-exposure. Furthermore, we estimate the lifetime of these broken states and the probability that exchange-competent configurations occur in them. We have also tested various sampling protocols and their ability to enhance the exploration of the broken states. Computational protocol used in this study can be applied to a broad range of sys-tems to gain valuable insight into the nature of the broken state, although further development is needed to devise a generally applicable quantitative method.

## 1 Introduction

In the folded state, the core of a protein is held together by hydrogen bond interactions involving backbone amides. Local fluctuations or complete unfolding of a protein breaks the hydrogen bonds involving backbone amides (*NH*) and allows water penetration. This enables hydrogen exchange (*HX*) between *NH* and the hydrogen atoms of the solvent water molecules. Thus, the *HX* rates, which can be measured by NMR spectroscopy or mass spectrometry, can give valuable information about the local stability of the protein structure as well as conformational changes and folding pathways [1].

Under typical conditions, NMR measurements yield the so-called *protection factor*, which can be expressed in terms of the observed exchange rate *k*_*HX*_:

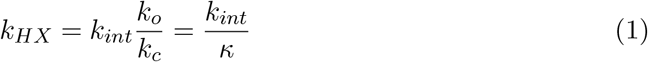

where *k*_*int*_ is the amide-specific intrinsic exchange rate of an unstructured peptide and *κ* is the protection factor which is defined as the ratio between the closing (*k*_*c*_) and opening (*k*_*o*_) rates [2, 3]. The free energy difference Δ*G*_*HX*_ between the exchange-competent (open) conformation and the dominant (closed) conformation can be expressed as:

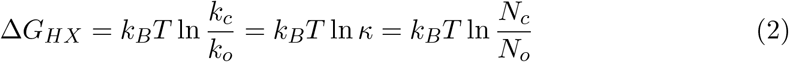

where *N*_*o*_ and *N*_*c*_ are fractional populations of open and closed states respectively [4, 2], *T* is the temperature and *k*_*B*_ is Boltzmann’s constant.

In principle, Δ*G*_*HX*_ for each residue can thus be determined directly by analysing a molecular dynamics (MD) trajectory and counting the number of configurations belonging to the open and closed state, respectively. However, this requires extensive sampling of the transitions between the closed and open states. In a recent study, a 250-microsecond trajectory of basic pancreatic trypsin inhibitor (BPTI) was analysed in this manner by *Persson and Halle* [4]. According to their classical HX model each amide exists either in a closed (C) state where there is no exchange or in an open (O) state where the exchange happens. The authors defined the O state as the configurations in which the amide hydrogen has at least two water oxygens within 2.6 Å and no other polar protein atom (except covalently neighbouring residues) within 2.6 Å. Hypothesising this geometrical criterion for the open state, the counting method gave a good agreement with experimental measurements of Δ*G*_*HX*_. The O states were found to have a lifetime of ∼100 ps.

Our hypothesis is that the O state, more precisely the exchange-competent configurations (ECC), is a rare fluctuation within a metastable ”broken” (B) state. The B state for a given amide is reached simply by breaking the backbone H-bond interaction, which typically will lead to influx of at least one water molecule close to the amide hydrogen. The B state is a metastable state and has a longer lifetime compared to the ECC. Figure 1 shows a pictorial representation of our proposed mechanism.

**Figure 1:**
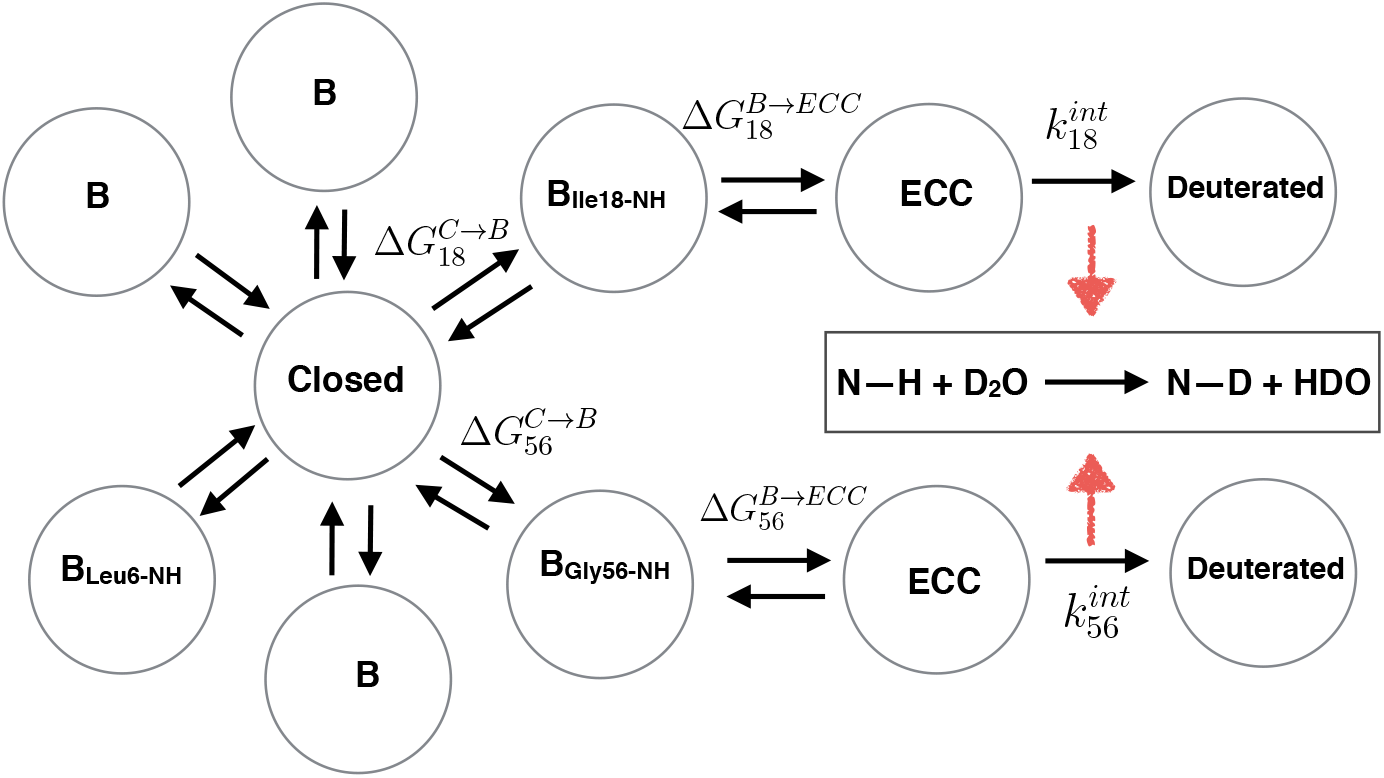
Pictorial representation of different conformational states of BPTI. In the closed state, the amide *NH* remain hydrogen bonded to other residues. The transition between closed and broken state can be sampled using MD or enhanced sampling methods, e.g. metadynamics.

In this project, we have tried to gain a deeper understanding of local fluctuations. As our test case, we have used BPTI, which is a small 53-residue protein that was used as a model system for comparing HX data with simulation results already 40 years ago [5]. We take a similar approach as ref. [4] in that we analyze the millisecond MD trajectory of the fully solvated native BPTI performed by Shaw et al. [6], which we denote *DESRES*. However, we extend the analysis and also perform enhanced-sampling simulations to try to sample the transitions in a more efficient way.

During the DESRES simulation, the *β* hairpin region of BPTI remained quite stable; hence the amides located in this region did not visit the open state. However, one amide (Ile18) located at the end of the *β* hairpin was able to access the open state due to local fluctuations involving a cooperative motion of neighbouring loops (residues 11 − −19 and 34 − −40) (Figure 2). This example illustrates the connection between general conformational changes in proteins and solvent rearrangements causing hydrogen exchange. In fact, this particular residue will be the main focus of the current study. However, we will also try to provide an overview of the dynamics of other residues and discuss the prospect of using enhanced-sampling methods to study them.

**Figure 2:**
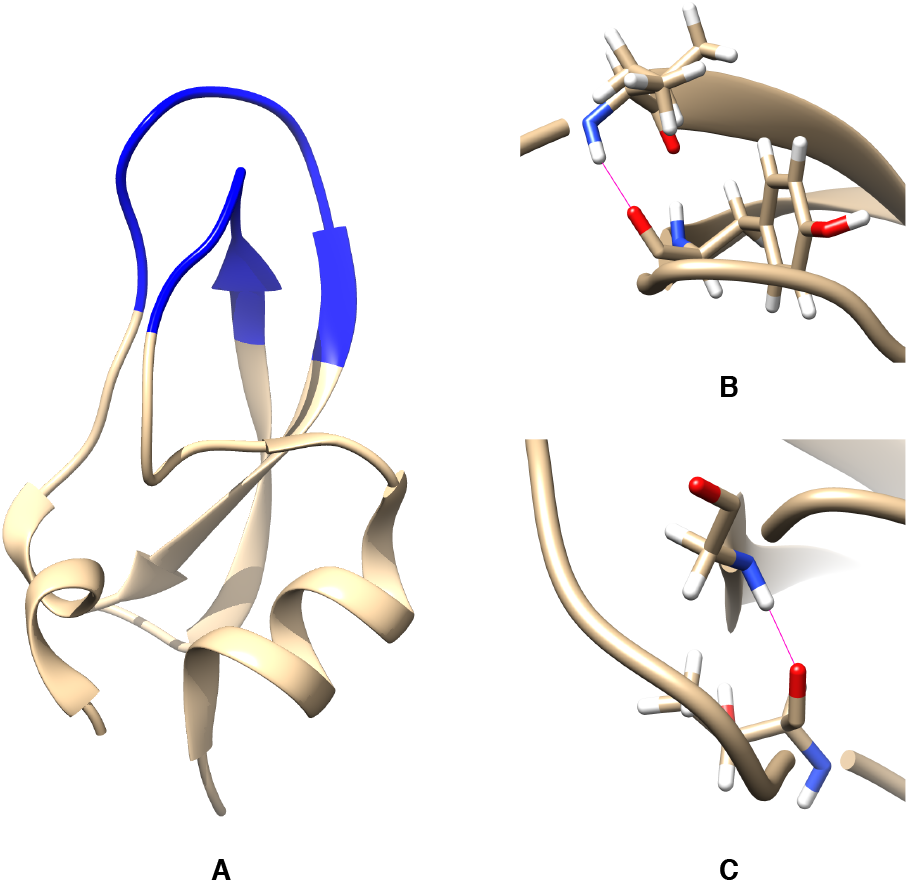
The loop involving residues 11-19 and 34-40 (highlighted in blue) in BPTI (A). The H-bonds (in magenta) involving Ile18-NH (B) and Gly36-NH (C) are also highlighted.

## 2 Methods

### 2.1 Definition of the M1 state

During the millisecond classical MD simulation, BPTI visited different long-lived states through rotation of the torsion angles around the Cys14–Cys38 disulphide bond. Previous investigations have identified five major states, M1, M2, M3, m_C38_, and m_C14_, out of which m_C14_ was most populated (50%) in the simulation [7]. On the other hand, experiments have shown that the M1 state dominates both in solution and in the crystal. Following [4], we therefore limited the study as much as possible to the M1 state, defined in Table 1. Thus, we used only the 1048369 frames (corresponding to 262 *μ*s of dynamics) of the DESRES trajectory that satisfy those criteria. For the new simulations, we tried two approaches: 1) to post-process the unrestrained trajectories by reweighting to the M1 state, and 2) to restrict the sampling to the M1 state. In the latter case, we used flat-bottomed restraining potentials (walls) as described in Table 1

**Table 1:** Criteria used to define the M1 state and to restrain new simulations to the M1 state by applying lower and upper harmonic wall potentials.

| Torsional angle | Min | Max | Lower wall | Upper wall | Force constant<br>(kJ mol<sup>-1</sup> deg<sup>-2</sup>) |
| --- | --- | --- | --- | --- | --- |
| $\chi_1$ (C14) | -120° | 0° | -100° | -20° | 0.5 |
| $\chi_2$ (C14) | -180° | 135° | 60° | 120° | 0.5 |
| $\chi_1$ (C38) | 0° | 120° | 40° | 100° | 0.5 |
| $\chi_2$ (C38) | | | -160° | -80° | 0.5 |
| $\chi_3$ (C14-C38) | 0° | 180° | 60° | 140° | 0.5 |

### 2.2 Analysis of hydrogen bond dynamics

Autocorrelation analysis of the hydrogen bonds was performed to estimate the lifetime of the broken states from the DESRES trajectory. For each hydrogen bond, the original distance time-series (sampled at 0.25 ns intervals) was first slightly smoothed using a running median of 5 frames. The purpose was to avoid single outliers that disturb the identication of initial points. Then, each frame was assigned a binary digit *m*(*t*) (0 for formed, 1 for broken hydrogen bond) depending on the distance being smaller or greater than a cutoff distance (typically 4 Å). The autocorrelation function *A*(*τ*) was then computed as follows:

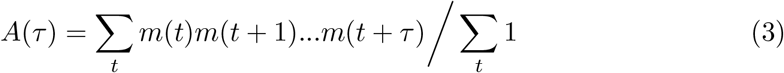

where the sum goes over all *initial points* of an interval of the broken state (i.e. satisfying *m*(*t*) = 1 and *m*(*t* − 1) = 0) and the purpose of the product is to ensure that only uninterrupted sequences of the broken state are counted. Special care was taken so that a time window did not include parts of separate trajectories or any large gaps created by the M1 filtering of the trajectory (although gaps of a total length of less than 20% of the window were accepted). The autocorrelation time was then calculated as the integral of the autocorrelation function:

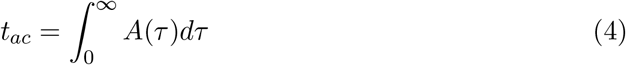

where in practice any upper integration limit significantly higher than the autocorrelation time could be used.

An additional lifetime analysis was performed by processing the original time series and keeping a running counter of how many of the last 20 frames had a *smaller* distance than the mean value of the whole series (typically around 3 Å). An interval of the broken state was then defined as an period during which the percentage of short-distance frames is constantly below a certain threshold, in our case chosen so low that no such frames were allowed at all. The idea is that, whereas there will always be fluctuations around any cutoff designed to separate broken from formed frames (creating difficult border cases), the probability of having a shorter-than-mean distance will be 50% in the formed state but close to zero in a truly broken state, thus providing a safe margin for avoiding falsely reported intervals. This procedure was indeed found to be more stable and was thus used to generate the histogram of lifetimes and to plot time-series along typical broken intervals. The histograms of lifetime distribution was generated by binning the list of intervals on a logarithmic scale, i.e. counting how many intervals that were of length 1 frame, 2 frames, 3–4 frames, 5–8 frames, 9–16 frames, etc.

### 2.3 Simulation setup

The system setup was kept as similar as possible to the DESRES simulation in order to compare results from DESRES. However, we used a more recent version of the Amber force field, *ff14sb* [8]. The water model was the same as used in DESRES, *TIP4P-Ew* [9]. Missing cysteine S–S bonds were added prior to simulation. The protein was solvated in a cubic box. The box size, number of water molecules and counter ions were kept identical as DESRES [6]. Prior to MD simulations, the structures were equilibrated for ∼ 3 ns in the NVT ensemble, where position restraints was applied on the *C*_*α*_ of the protein with a harmonic restraint potential with a force constant of 500 kJ mol^−1^ nm^−2^. Long-range electrostatic interactions were treated using the particle mesh Ewald (PME) method [10] with a Fourier spacing of 0.16 nm and a cut-off for van der Waals and short range electrostatics of 1.2 nm. Finally, production runs of varied lengths were carried out in the NVT ensemble using a time-step of 2 fs. The bonds of all hydrogen atoms were constrained using the LINCS algorithm. and the temperature was kept at 300 K using a velocity rescaling thermostat [11].

All simulations were performed using Gromacs 2018 [12] patched with Plumed 2.4 [13].

### 2.4 tICA analysis

The M1 subset [7] of the DESRES trajectory was used to perform time-lagged independent component analysis (tICA) using using *MSM Builder 3*.*8* [14]. Before performing tICA, rotational and translational degrees of freedom were removed from the trajectory. tICA was performed by transforming the raw *all-atom* cartesian co-ordinates into a subset of the *C*_*α*_ co-ordinates for residues 6 − −56 using *Rawpositionfeaturiser*. The *lag* time was set to 100 ns. The first five independent components (e.g. tIC1, tIC2, tIC3 etc.) were used for subsequent analyses. tIC coefficients corresponding to each *C*_*α*_ atom were extracted using an in-house *Python* script. *Plumed driver* [13] was used to extract projections along independent components from the DESRES trajectory. Independent components generated by tICA were subsequently used in enhanced-sampling simulations.

### 2.5 Metadynamics

Metadynamics simulations were performed using distance and tIC1 (first time-independent component) as collective variables. tIC1 was used within a parallel tempering metady-namics workflow as described below.

### 2.5.1 Distance-CV

We have used the Ile18-NH–O-Tyr35 H-bond distance (N..O distance) as collective variable to perform well-tempered metadynamics simulations. Metadynamics simulations were performed using Gaussian height 1.2 kJ/mol, width 0.005 nm and bias factor 15. Upper and lower harmonic wall potentials were used to keep the system within the M1 state as described above. Additional upper and lower walls at 0.6 nm and 0.2 nm respectively, with a (harmonic) force constant of 5000 kJ mol^−1^ nm^−2^, were placed along the H-bond distance to restrict the sampling from regions of high energy. Furthermore, we have performed metadynamics simulations using H-bond distance involving Met52-NH (Met52-NH–O-Ala48)^1^ and Gly36-NH (Gly36-NH–O-Thr11)^2^ using identical Gaussian height, width and bias factor.

### 2.6 Parallel tempering metadynamics

In parallel-tempering [15] (also called replica exchange), replicas of the system are simulated using the same force-field but at different temperatures. Frequent exchanges with higher temperature replicas prevents the normal-temperature replica from getting trapped in local minima. Parallel tempering combined with metadynamics significantly enhances the sampling of the conformational space. We performed parallel-tempering (PT) in the well-tempered ensemble (WTE) using 16 replicas spanning a temperature range of 300 K to 432 K. In WTE [16], bias was applied on the systemś potential energy using Gaussian height 1.2 kJ/mol and width 140 kJ/mol, with a bias factor of 9. Before starting to apply bias on tIC1, a pure PT-WTE simulation was carried out for 18 ns, while building up the WTE bias potential. The end-point of this simulation was used to perform PT-WTE-metadynamics (denoted PT-METAD) using tIC1 as CV with Gaussian height 0.5 kJ/mol, width 0.00005 and bias factor 10. The WTE bias potential was kept fixed during this simulation and replica exchange was attempted every 2 ps.

### 2.7 Reweighting

Reweighting allows one to remove the effect of bias from the time-dependent metadynamics potential and external wall potentials, and recover the unbiased probability distribution along any reaction co-ordinate. In this study, we have used the reweighting algorithm proposed by Tiwary and Parrinello [17]. The unbiased probability distribution was collected in one- or two-dimensional histograms using Gaussian kernels with recommended bandwidth and grid spacing.

### 2.8 Short MD simulations

Several short MD simulations were performed starting with broken conformations of Ile18-NH. Arbitrary conformations with broken hydrogen bond were dumped from different snapshots of a metadynamics simulation with distance CV. Because the bias potential was suddenly turned off to start MD simulations, a new restrained equilibration was performed for each simulation for ∼ 3 *ns* with the same protocol as the initial equilibration.

## 3 Results and Discussion

Using the definition of exchange-competent (”open”) configurations from Ref. [4], it was previously found that the lifetimes of these configurations were short for all amides (81 ± 18 ps). In fact, as shown in Fig. 3, the maximum number of consecutive exchange-competent frames in the DESRES data is consistent with a random distribution of these frames along the simulation.

**Figure 3:**
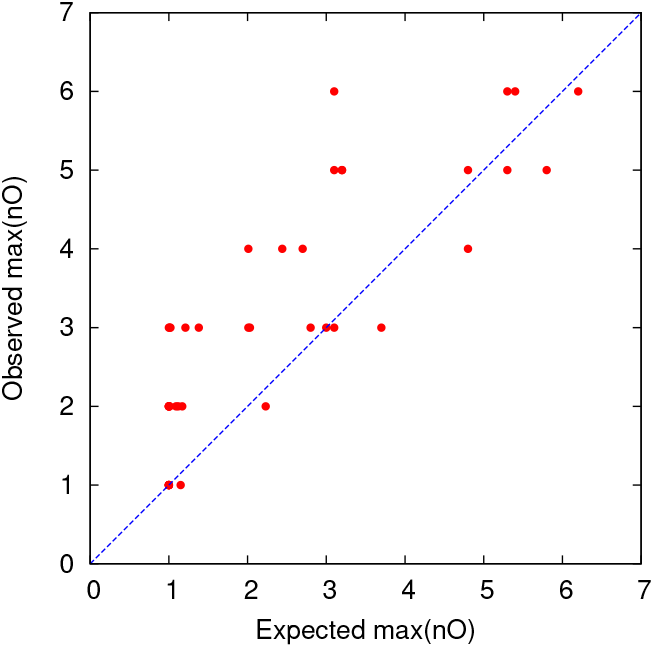
The observed maximum number of consecutive ”open frames” in DESRES [4] plotted against the expected number if the same number of open frames were completely randomly distributed over the trajectory. Each point corresponds to one residue and the line indicates perfect agreement.

For many of the hydrogen bonds, the lifetime of the “broken” state is much longer, as discussed further below. A plausible hypothesis is therefore that the metastable state corresponding to a solvent-exposure of a given amide hydrogen is in fact the broken state, whereas the actual ”exchange-competent” configurations with two water molecules are rare but regular occurrencies within this state, corresponding to the outermost tail of the water distribution. The aim of the current study is to test this hypothesis for most of the residues, and to investigate these states in more detail for a small subset of residues.

### 3.1 Lifetime of the broken state

#### 3.1.1 Analysis of the DESRES trajectory

A common way to define lifetimes of e.g. hydrogen bonds is to analyze the autocorrelation function of a step function (bond formed or not) [18]. For the lifetime of the broken state, we performed an analogous analysis, as described in the method section. The calculated autocorrelation time for the most relevant hydrogen bonds are shown in Table 2 as a function of the cutoff distance (the rest are either very strong or very weak and thus of less interest). As can be seen, the results are rather insensitive to the cutoff distance in some interval around 4 Å, with the exception of the Met52–Ala48 bond, and thus a cutoff distance of 4 Å was used in the following. The corresponding autocorrelation functions are shown in Fig. 4.

**Table 2:** Autocorrelation time for the broken state of various hydrogen bonds from the DESRES simulation, using three different cutoff distances.

| Bond | Autocorrelation time (ns) |  |  |
| --- | --- | --- | --- |
|  | 3.5 Å | 4.0 Å | 4.5 Å |
| Glu7–Phe4 | 24.1 | 26.2 | 17.4 |
| Ile18–Tyr35 | 0.8 | 0.9 | 0.7 |
| Asn24–Leu29 | 6.4 | 16.2 | 17.4 |
| Gly28–Asn24 | 16.9 | 17.1 | 17.5 |
| Gly36–Thr11 | 4.7 | 3.9 | 3.6 |
| Cys51–Ser47 | 1.8 | 0.7 | 0.5 |
| Met52–Ala48 | 33.0 | 11.2 | 1.9 |
| Arg53–Glu49 | 45.4 | 41.8 | 45.5 |
| Cys55–Cys51 | 61.0 | 69.3 | 75.1 |

**Figure 4:**
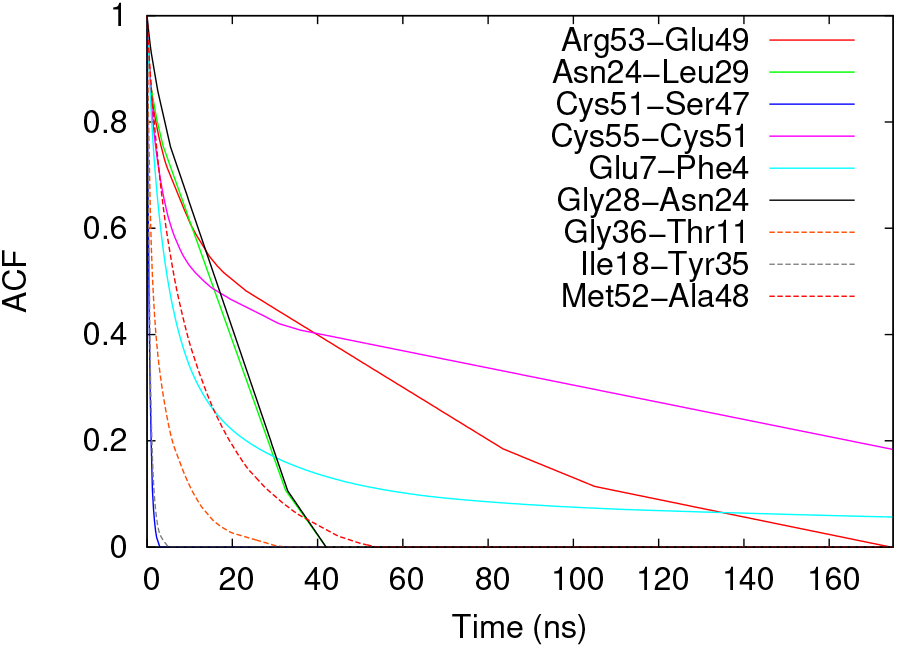
Autocorrelation function of the broken state for various hydrogen bonds from the DESRES simulation using a cutoff distance of 4.0 Å.

The distributions of lifetimes for each broken state is shown in Fig. 5. Clearly, the individual visits to the broken state vary substantially in length, but for all the shown bonds, more than half of the visits are shorter than 10 ns and for all but one bond (Ile18–Tyr35), more than half of the visits are longer than 2 ns, so the range 2–10 ns can be considered quite typical for the duration of visits to the metastable broken state. This timescale is significantly longer than the lifetime of the exchange-competent (“open”) state described in Ref. [4].

**Figure 5:**
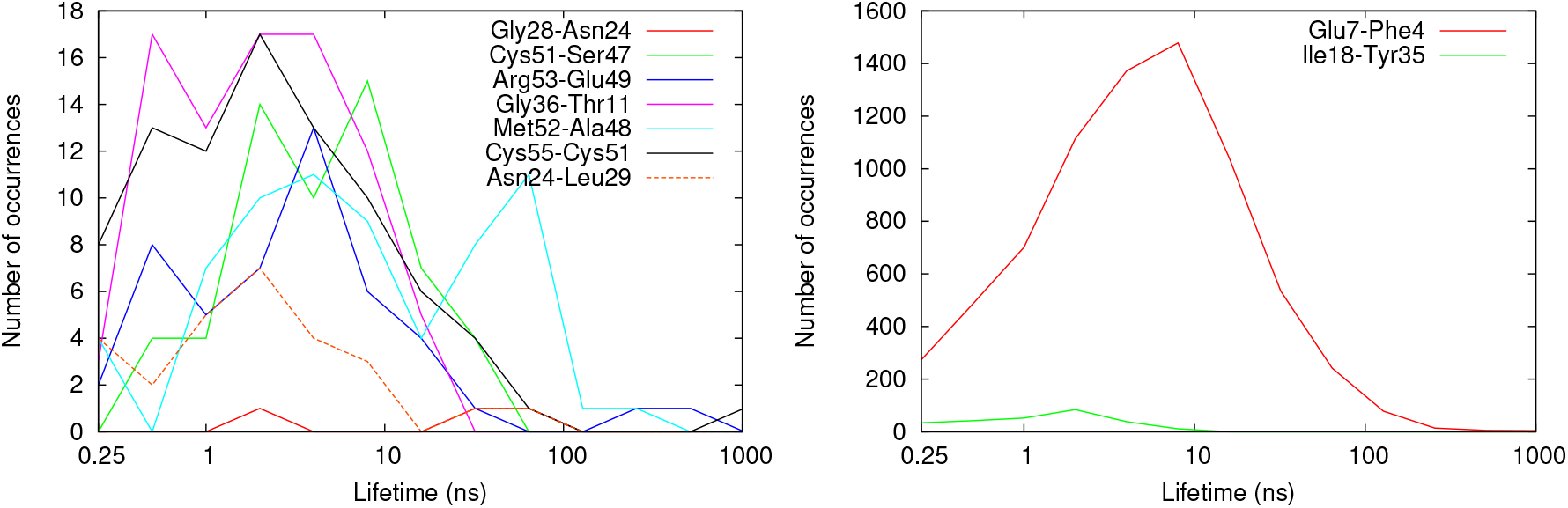
Histogram showing the distribution of lifetimes for the broken state of various hydrogen bonds (note the logarithmic scale on the x axis). The actual number of occurences of a broken state with a lifetime in a given interval is shown; the total number of such occurrences varies between 3 and 7342 for the various hydrogen bonds.

#### 3.1.2 New MD simulations

Some of the broken states have so short lifetime that the accuracy is severely compromised by the sparsity of the DESRES trajectory (frames saved every 250 ps). Therefore, for the Ile18–Tyr35 hydrogen bond, we ran additional MD simulations starting from the broken state and monitored the hydrogen-bond distance until the hydrogen bond was reformed. As described in the method section, the initial coordinates were taken from a biased ensemble (generated through metadynamics on the hydrogen-bond distance), so some caution is needed in the interpretation, because the force acting to reform the hydrogen bond might be stronger than that occurring in an equilibrium ensemble; however, no indication of unphysical behavior was observed. The autocorrelation time for 16 combined trajectories with different initial structures was 0.67 ns (using a cutoff distance of 4.0 Å), in good agreement with Table 2. Each simulation actually showed a clear transition, occurring 1.8 ± 1.4 ns into the unconstrained simulation on average (approximately following a Poisson distribution). A more detailed view of these transitions will be shown below.

### 3.2 Solvent-exposure in the broken state

Next, we investigated how the solvent-exposure of the amide differs between the broken state and the closed state. The distribution of first- and second-water distances for the ensembles with formed and broken hydrogen bond, respectively, are shown in Fig. 6. For most of the amides, there is a clear correlation between the breakage of the hydrogen bond and the increased solvent-exposure. For example, the most probable distance from the Cys55 amide hydrogen to the closest water molecule is reduced from 4.9 to 2.0 Å upon breaking the intra-helical hydrogen bond to the oxygen of Cys51.

**Figure 6:**
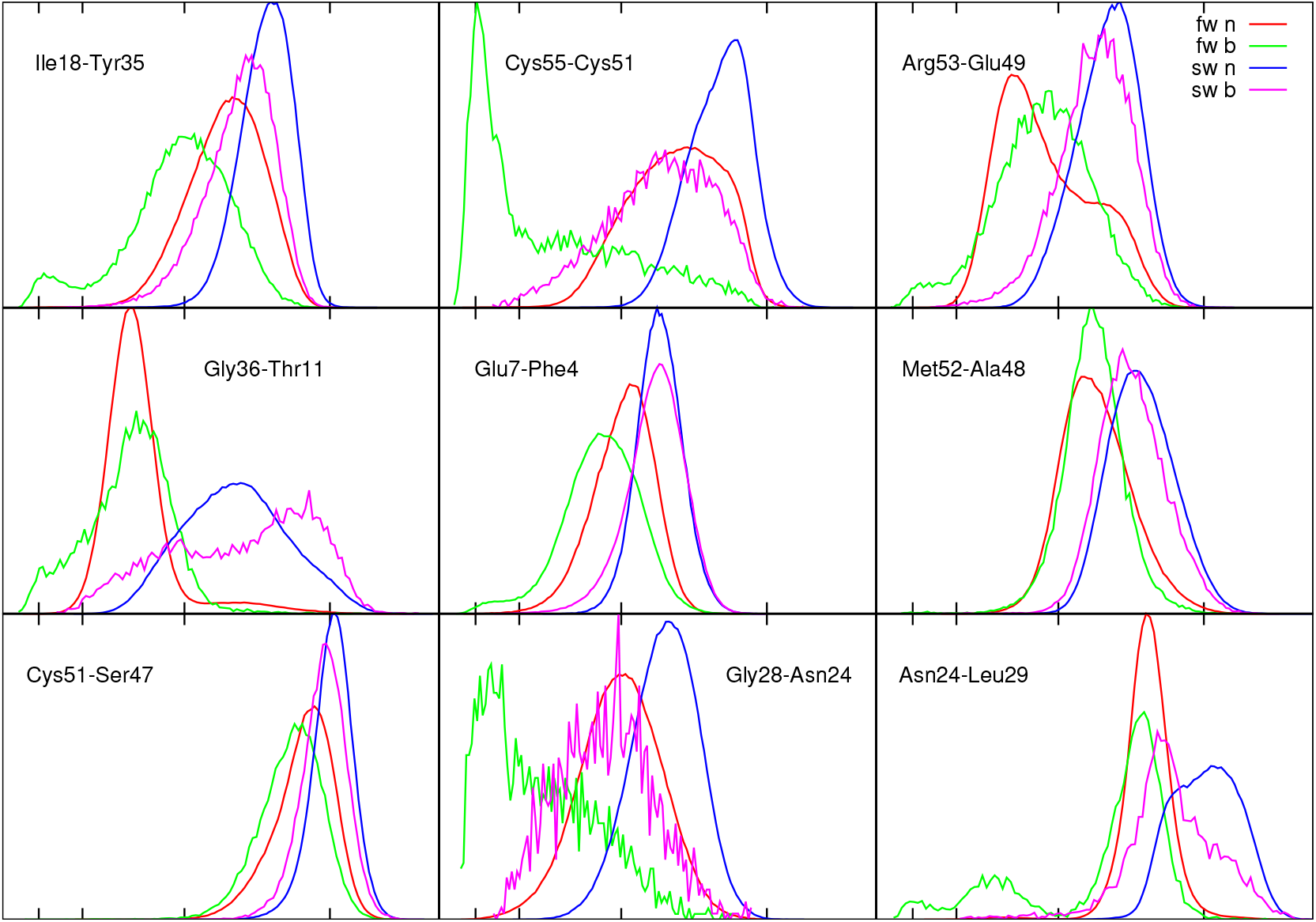
Histograms showing the distribution of the distance from the amide hydrogen to the first (fw) and second (sw) closest water oxygen, respectively, in the normal (n) and broken (b) state, respectively. The states are defined as having a N..O hydrogen bond length smaller or larger than 3.5 Å. The tics on the x axis marks distances of 2 Å, 2.6 Å, 4 Å, and 6 Å. The line colors shown in the upper right diagram apply to all diagrams (red = first water in normal state, green = first water in broken state, blue = second water in normal state, magenta = second water in broken state).

The distributions of the SW distance allows us to estimate the frequency of exchangecompetent configurations (ECCs; denoted as ”open” configurations in Ref. [4]) occurring in the broken state, which can be converted to an apparent free energy 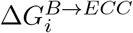(see Figure 1) through the following relation:

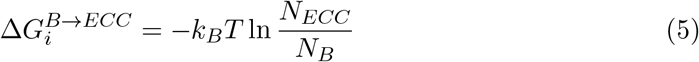

where *N*_*ECC*_ is the number of exchange-competent frames (i.e. broken and with *SW <* 0.26 nm) and *N*_*B*_ is the number of broken frames. It is important to note that we do not have to assume ECC to be a metastable state to use this equation. The results are given in Table 3. Interestingly, there is a great variation between various residues, which probably reflects the variation in geometry and specific interactions occuring in the surroundings of each amide.

**Table 3:** Apparent free energy (in kJ/mol) of accessing exchange-competent configurations when already in the broken state, calculated using Eq. 5 based on the DESRES and PT-METAD data, respectively. The number of ECC frames that each calculation is based on is also shown. Only entries with *N*_*ECC*_ ≥ 5 are included.

| Amide | DESRES |  | PT-METAD |  |
| --- | --- | --- | --- | --- |
| | $\Delta G_i^{B \rightarrow ECC}$ | $N_{ECC}$ | $\Delta G_i^{B \rightarrow ECC}$ | $N_{ECC}$ |
| Cys-5 | 17.9 | 144 |  |  |
| Leu-6 | 14.4 | 711 |  |  |
| Glu-7 | 19.2 | 283 |  |  |
| Ile-18 | 19.2 | 27 | 16.5 | 61 |
| Gly-28 | 9.3 | 39 | 5.7 | 29 |
| Leu-29 | 28.9 | 5 |  |  |
| Gly-36 | 12.4 | 102 | 12.6 | 294 |
| Met-52 | 22.3 | 5 |  |  |
| Arg-53 | 16.2 | 46 |  |  |
| Thr-54 | 18.2 | 167 |  |  |
| Cys-55 | 12.7 | 58 |  |  |
| Gly-56 | 10.9 | 2012 | 16.3 | 22 |

It is also interesting to know whether the increased solvent-exposure coincides exactly with the broken hydrogen bond or whether there are time lags. The time variation of the hydrogen bond length and the distances to the closest and second closest water molecules is shown in Fig. 7 for six hydrogen bonds (those showing highest correlation) during a time interval encompassing the most long-lived occurrence of the broken state (results for other long-lived occurrences were visually confirmed to be similar but are not shown). In most cases, the decrease in first-water distance occurs almost instantaneously when the hydrogen bond is broken. During some broken state periods, there are several subperiods with varying solvation behavior.

**Figure 7:**
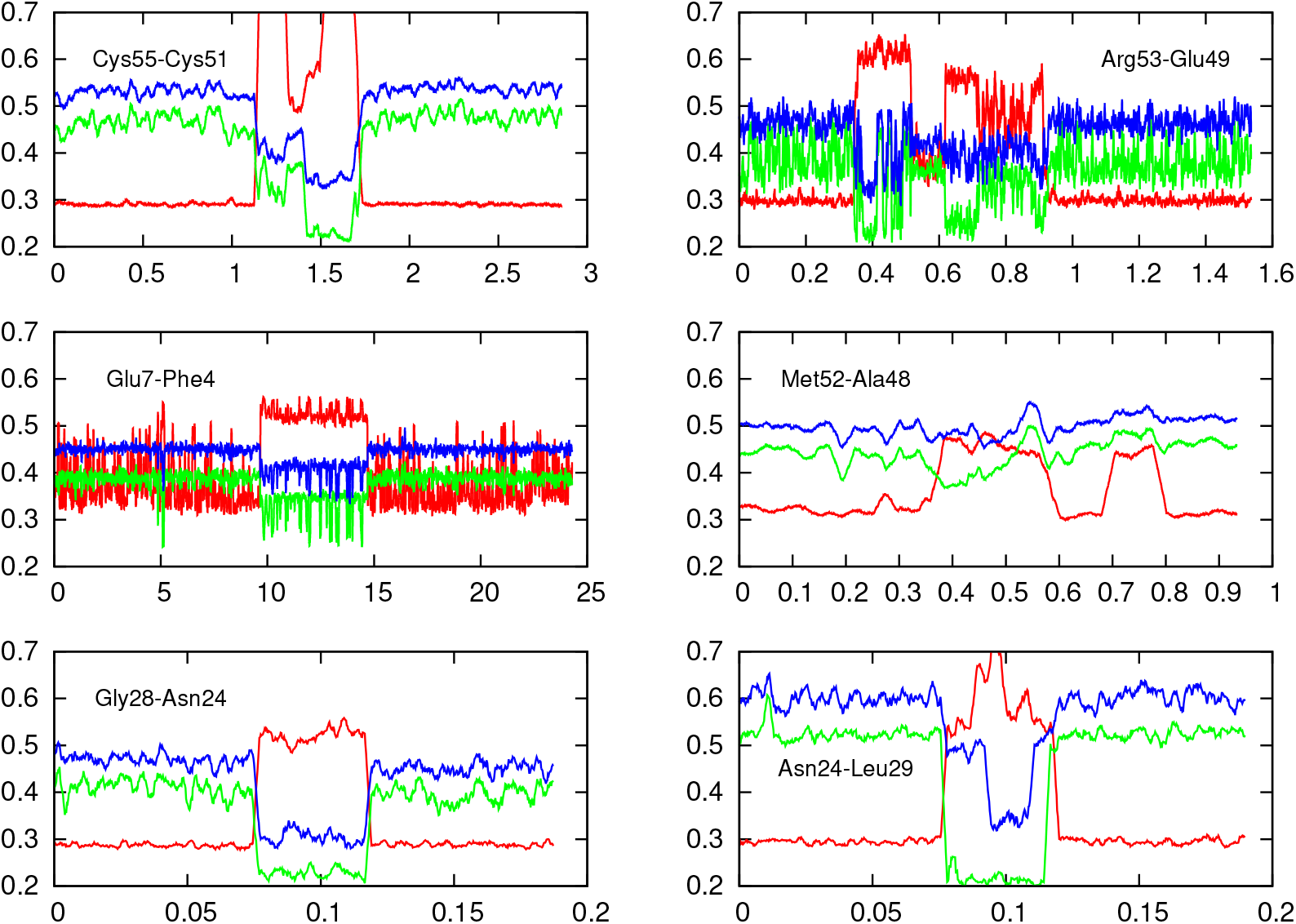
Hydrogen bond length (red) and the distances between the amide hydrogen and the first (green) and second (blue) water molecule, respectively, as a function of time (in microseconds), around the most long-lived occurrence of the broken state in the DESRES simulation. For each hydrogen bond, the starting time of the displayed time interval has been set to zero; the absolute starting times are 15.94, 16.73, 150.36, 148.92, 768.28 and 768.28 *μ*s into the MD simulation, respectively

The previously mentioned short MD simulations of the reforming of the Ile18–Tyr35 hydrogen bond have the advantage of more densely saved trajectories and thus allows us to get a more detailed view of the actual transition. Figure 8 shows the time evolution of the hydrogen bond distance together with the FW and SW distances around the transition, both for all 16 trajectories averaged and for a specific example trajectory. As can be seen, the transition occurs at a timescale of a few hundred picoseconds, with the hydrogen bond distance gradually decreasing simultaneously as the water molecules get closer. Histograms of the distributions of these distances during the short MD simulations are shown in Figure 9. Compared to the corresponding diagram in Fig. 6, we see that the distributions are much more separated between the states, due to the more clean-cut transitions.

**Figure 8:**
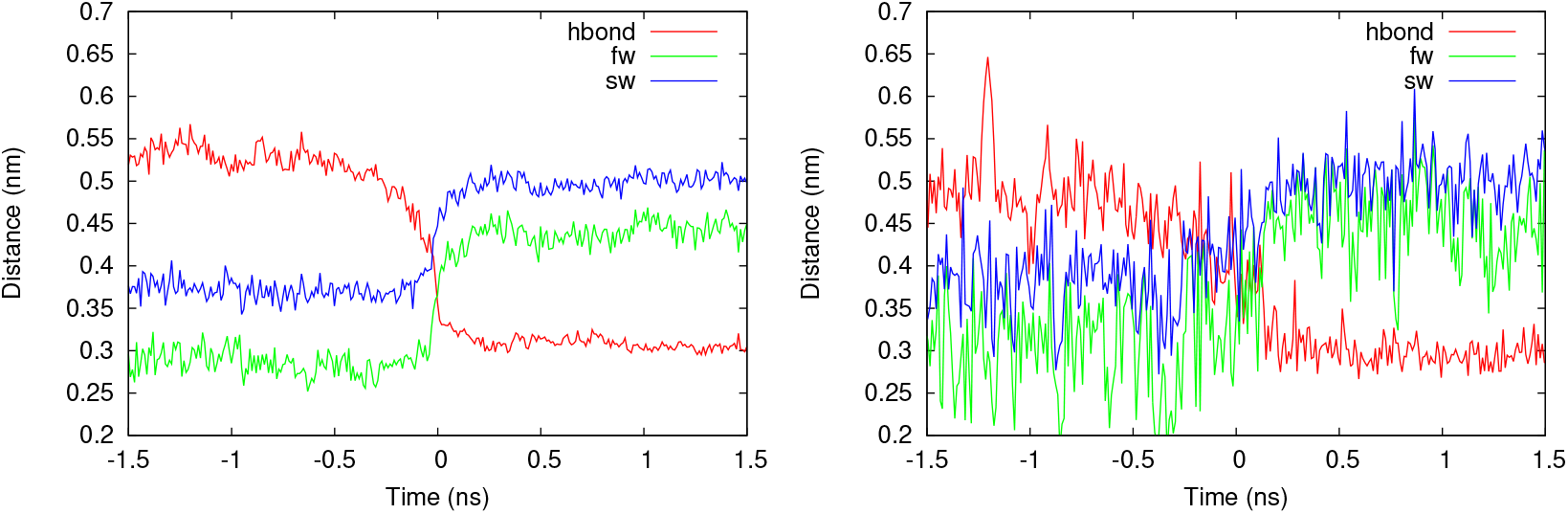
Hydrogen bond length and the distances between the amide hydrogen and the closest and second closest water molecule, respectively, as a function of time, around the reformation of the Ile18–Tyr35 hydrogen bond in MD simulations started from the broken state. The left panel shows the average over all 16 trajectories (after temporal alignment of the transition). The right panel shows one particular example trajectory

**Figure 9:**
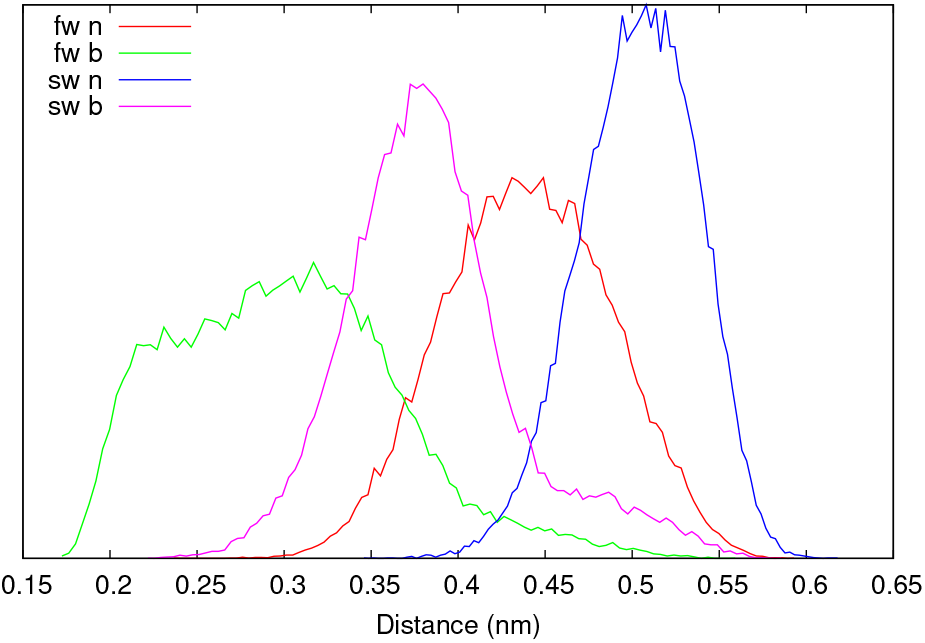
Distribution of first water (fw) and second water (sw) distances in the normal (n) and broken (b) states during the short MD simulations started from the broken state of the Ile18–Tyr35 hydrogen bond.

### 3.3 Free-energy landscape for breaking hydrogen bonds

To gain a better understanding of the energetics of hydrogen bond breaking and amide solvent exposure, we performed enhanced-sampling simulations using several different sets of collective variables (CVs). Enhanced sampling allows us to collect more statistics in a shorter time than regular MD simulations, and can therefore provide more reliable results. The dynamics, on the other hand, is less straightforward to recover from a biased simulation and thus will not be further discussed.

There is always a risk that the results are influenced by the choice of CVs. Therefore, we tried to obtain as much information from a simulation with “automatically” devised CVs (section 3.3.1). Additionally, we performed simulations using CVs that were es-pecially designed to promote the breaking of specific hydrogen bonds (section 3.3.2). We paid particular attention to cross-validation of the results obtained with the general and more specific CVs, respectively, and, when possible, also with the results from the DESRES simulation.

#### 3.3.1 PT-METAD simulation based on tICA analysis

Time-lagged independent component analysis (tICA) provides a way to analyze an MD trajectory and extract a few linear vectors describing the slowest processes in the system, which can then be used as CVs in enhanced-sampling simulations to help the exploration of these processes. In our case, we used the DESRES trajectory as input, and a procedure described in the method section, to extract the primary such component (denoted tIC1), as well as additional components which were mainly used for testing and will not be further discussed.

The maxima of local backbone flexibility are found in the regions of residues 11–19 and 34–40 (and in the terminal residues). This was also confirmed by visual inspection of the DESRES trajectory. Therefore, it was not surprising that tIC1 mainly involves these loops. For example, a scatter plot of tIC1 vs tIC4 colored by the C_*α*_ RMSD of the loop region (towards the crystal structure after alignment on all C_*α*_ atoms) showed that the well separated “right region” (tIC1*>* 0.0028) captured the higher RMSD of the loop residues (Figure 10 A). It is interesting to note that the population of broken H-bonds (*>* 0.37 nm) for Ile18 and Gly36 is higher on the right side. As a broken H-bond leads to influx of water, a corresponding pattern is seen for the FW distance for both the Ile18 and Gly36 amides (Figure 10).

**Figure 10:**
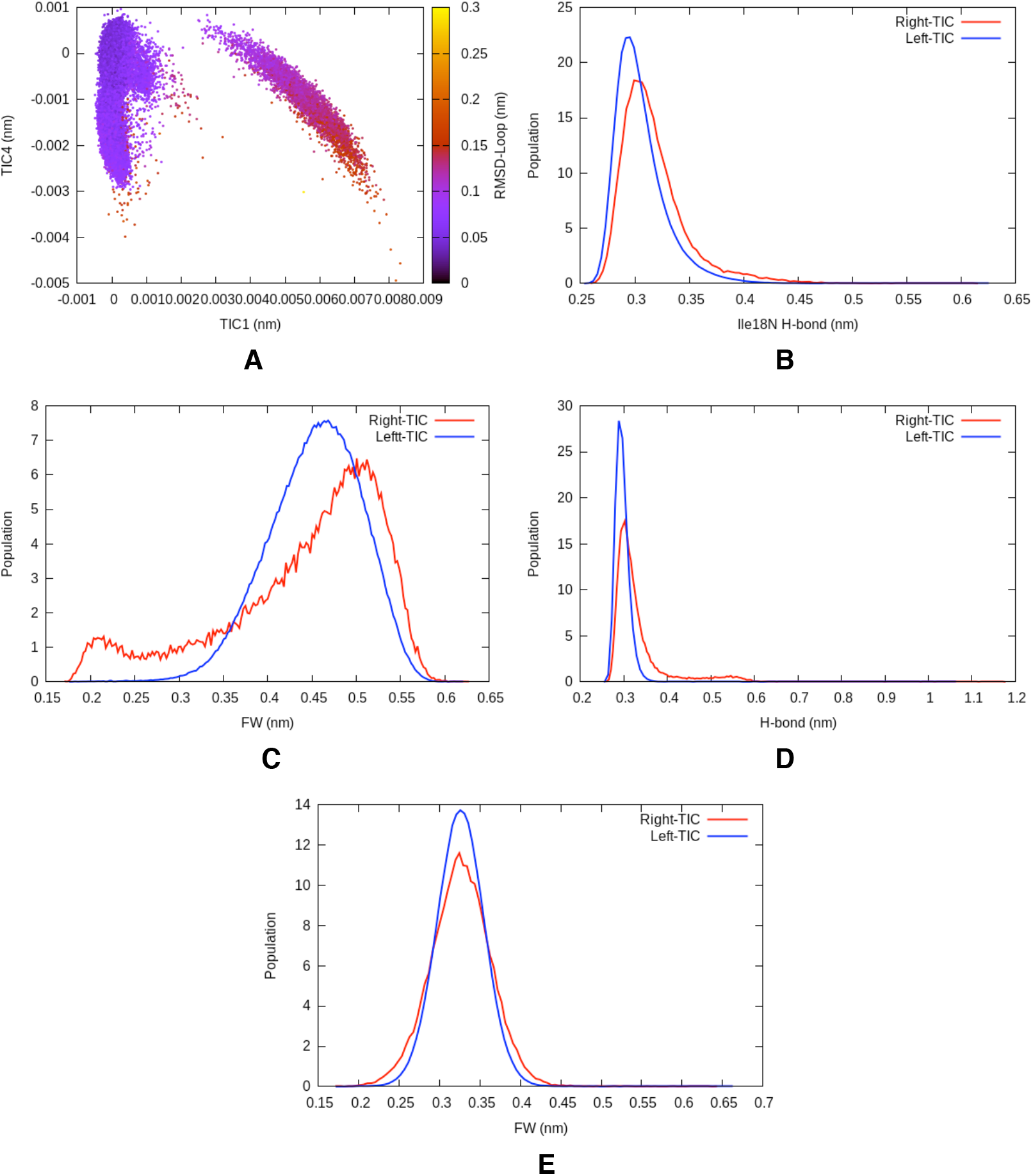
tIC1 vs tIC4 plotted against RMSD of the loop residues during DESRES-M1 (A). 1D histogram showing population of Ile18 and Gly36 H-bond distance within right (tIC1*>*0.0028 nm) and left (tIC1<0.0028 nm) region (B and D respectively). 1D histogram showing distance of FW within right and left tIC1 for Ile18 and Gly36 amides (C and E respectively).

Preliminary testing showed that metadynamics using only tIC1 as CV suffered from convergence problems, and that addition of further CVs, such as tIC2 or tIC4, did not significantly improve the situation. On the other hand, raising the temperature improved the convergence significantly. Therefore, we settled for a combination of parallel tempering and metadynamics (denoted PT-METAD), in which only tIC1 was specifically biased, but where the parallel tempering allowed overcoming kinetic barriers in other slow degrees of freedom of the system.

A PT-METAD simulation with 16 replicas spanning a temperature range of 300– 432 K was setup as described in the method section and run for 230 ns (i.e. in total 3.7 *μ*s for all the replicas) with restraints keeping it within the M1 state. For validation, an identical PT-METAD simulation without restraints was also performed, in which the restriction to M1 was done through a postprocessing step, but only the restrained simulation will be discussed here, as the results were very similar.

The most relevant connection to experimental HX data is the solvent exposure of the amides. Table 4 shows the percentage of the simulation time spent in various “solvent-exposed” configurations, both in the PT-METAD simulation (the 300 K replica) and in the DESRES simulation for reference. Significantly increased sampling of solvent-exposed configurations were observed for Ala-16, Ile-18, Arg-20, Tyr-35, Gly-36, and Gly-37, whereas the rest of the amides showed rather similar percentages as in DESRES. This set of affected residues is consistent with an enhanced fluctuation of the loops that they are situated in. It is important to note that the “raw” percentage is only used as an indicator of whether the sampling is enhanced or not, and does not correspond to any observable, because the ensemble is biased. However, the unbiased percentage can be recovered from the biased ensemble through a reweighting algorithm. The reweighted results are also shown in Table 4, and it’s encouraging to see that reweighting brings the increased percentages back to a level similar to the DESRES results.

**Table 4:** Percentage of the trajectory frames satisfying A) the criterion FW< 0.25 nm or B) the ”open-state” criterion of Ref. [4], SW< 0.26 nm. For each criterion, data for the DESRES ensemble (DES) and the raw PT-METAD ensemble (PT) are given, as well as the ratio between PT-METAD and DESRES. In addition, the reweighted percentage from the PT-METAD ensemble (i.e. eliminating the effect of the bias potential) is given (PT-R). The rows are arranged so that the residues showing significantly enhanced sampling in the PT-METAD simulation are listed above the dividing line and the rest are listed below the line for reference (21, 22, 23, 33, 45, 51 and 52 were omitted because of uninterestingly small values).

| Residue | Criterion A ($FW < 0.25$ nm) | | | | Criterion B ($SW < 0.26$ nm) | | | |
| --- | --- | --- | --- | --- | --- | --- | --- | --- |
|  | DES | PT | PT-R | ratio | DES | PT | PT-R | ratio |
| Ala16 | 14.66 | 67.69 | 23.10 | 4.6 | 0.23 | 2.12 | 0.48 | 9.1 |
| Ile18 | 0.41 | 12.81 | 0.43 | 31.4 | 0.00 | 0.03 | 0.00 | 9.9 |
| Arg20 | 0.01 | 0.30 | 0.01 | 29.3 | 0.00 | 0.00 | 0.00 |  |
| Tyr35 | 0.03 | 1.05 | 0.01 | 30.0 | 0.00 | 0.00 | 0.00 | 18.0 |
| Gly36 | 0.66 | 5.93 | 0.51 | 9.0 | 0.01 | 0.13 | 0.00 | 11.9 |
| Gly37 | 2.26 | 33.50 | 7.37 | 14.8 | 0.01 | 0.25 | 0.00 | 33.1 |
| Cys38 | 81.82 | 82.01 | 91.02 | 1.0 | 0.01 | 0.07 | 0.00 | 6.5 |
| Asp3 | 82.07 | 77.84 | 77.92 | 0.9 | 8.28 | 7.82 | 8.39 | 0.9 |
| Phe4 | 12.92 | 17.01 | 16.83 | 1.3 | 0.05 | 0.02 | 0.02 | 0.3 |
| Cys5 | 3.22 | 0.02 | 0.01 | 0.0 | 0.01 | 0.00 | 0.00 | 0.0 |
| Leu6 | 3.15 | 0.32 | 0.19 | 0.1 | 0.07 | 0.00 | 0.00 | 0.1 |
| Glu7 | 1.70 | 0.30 | 0.21 | 0.2 | 0.03 | 0.00 | 0.00 | 0.0 |
| Tyr10 | 80.87 | 79.37 | 83.29 | 1.0 | 0.35 | 0.49 | 0.37 | 1.4 |
| Thr11 | 81.45 | 83.96 | 83.52 | 1.0 | 9.67 | 13.65 | 13.49 | 1.4 |
| Gly12 | 66.55 | 70.13 | 72.33 | 1.1 | 0.66 | 0.69 | 0.93 | 1.0 |
| Cys14 | 49.99 | 46.17 | 59.21 | 0.9 | 0.00 | 0.01 | 0.00 |  |
| Lys15 | 87.36 | 83.77 | 88.26 | 1.0 | 6.93 | 4.26 | 6.66 | 0.6 |
| Arg17 | 65.83 | 34.29 | 58.92 | 0.5 | 2.03 | 0.85 | 2.15 | 0.4 |
| Ile19 | 82.31 | 78.31 | 79.93 | 1.0 | 0.96 | 0.89 | 0.86 | 0.9 |
| Asn24 | 0.04 | 0.08 | 0.00 | 1.8 | 0.00 | 0.00 | 0.00 |  |
| Ala25 | 80.15 | 78.83 | 76.81 | 1.0 | 10.31 | 13.11 | 12.27 | 1.3 |
| Lys26 | 24.62 | 30.65 | 31.30 | 1.2 | 0.04 | 0.04 | 0.04 | 0.8 |
| Ala27 | 0.14 | 0.09 | 0.08 | 0.7 | 0.00 | 0.00 | 0.00 |  |
| Gly28 | 0.65 | 0.17 | 0.12 | 0.3 | 0.00 | 0.01 | 0.00 | 2.8 |
| Leu29 | 0.02 | 0.04 | 0.00 | 2.3 | 0.00 | 0.00 | 0.00 | 0.0 |
| Cys30 | 72.13 | 80.26 | 78.63 | 1.1 | 0.93 | 1.40 | 1.43 | 1.5 |
| Gln31 | 0.08 | 0.04 | 0.07 | 0.5 | 0.00 | 0.00 | 0.00 |  |
| Thr32 | 73.26 | 76.33 | 76.63 | 1.0 | 3.32 | 3.42 | 3.73 | 1.0 |
| Val34 | 74.10 | 76.90 | 79.61 | 1.0 | 1.45 | 1.45 | 1.65 | 1.0 |
| Arg39 | 88.34 | 90.47 | 90.23 | 1.0 | 9.42 | 10.01 | 11.44 | 1.1 |
| Ala40 | 63.33 | 70.29 | 63.99 | 1.1 | 0.78 | 1.26 | 0.60 | 1.6 |
| Lys41 | 93.92 | 91.31 | 95.73 | 1.0 | 0.03 | 0.16 | 0.00 | 5.5 |
| Arg42 | 56.41 | 78.23 | 76.13 | 1.4 | 2.29 | 3.32 | 2.98 | 1.5 |
| Asn43 | 67.43 | 78.91 | 79.28 | 1.2 | 1.96 | 2.46 | 2.72 | 1.3 |
| Asn44 | 0.88 | 1.10 | 0.40 | 1.2 | 0.00 | 0.01 | 0.00 | 4.1 |
| Lys46 | 68.27 | 94.61 | 95.15 | 1.4 | 1.98 | 4.32 | 4.26 | 2.2 |
| Ser47 | 1.97 | 4.56 | 4.46 | 2.3 | 0.00 | 0.01 | 0.00 | 1.5 |
| Ala48 | 84.15 | 87.06 | 87.23 | 1.0 | 3.01 | 1.57 | 1.92 | 0.5 |
| Glu49 | 40.94 | 18.62 | 18.76 | 0.5 | 0.03 | 0.01 | 0.02 | 0.2 |
| Asp50 | 0.10 | 0.04 | 0.05 | 0.4 | 0.00 | 0.00 | 0.00 |  |
| Arg53 | 0.19 | 0.09 | 0.11 | 0.5 | 0.00 | 0.00 | 0.00 | 0.0 |
| Thr54 | 0.87 | 0.01 | 0.00 | 0.0 | 0.02 | 0.00 | 0.00 | 0.0 |
| Cys55 | 0.46 | 0.00 | 0.00 | 0.0 | 0.01 | 0.00 | 0.00 | 0.0 |
| Gly56 | 9.51 | 3.12 | 2.82 | 0.3 | 0.24 | 0.02 | 0.02 | 0.1 |
| Gly57 | 74.58 | 74.29 | 72.81 | 1.0 | 8.04 | 7.84 | 7.48 | 1.0 |
| Ala58 | 68.66 | 71.89 | 72.89 | 1.0 | 6.83 | 6.57 | 7.13 | 1.0 |

We chose two of the residues that displayed an enhanced solvation, Ile-18 and Gly-36, and performed a more detailed analysis.

### Ile-18

The DESRES simulation did a poor sampling of the broken state of Ile18, especially when only the M1 state was considered. PT-Metad did a much better sampling and several recrossings were observed along tIC1 as well as the Ile18–Tyr35 hydrogen bond. Convergence was monitored in terms of the free-energy difference between basins in the one-dimensional free-energy profiles with respect to tIC1, the hydrogen bond distance, and the FW distance, respectively (Figure 11). All three curves converge after ∼ 150 ns but to different values. This might seem counter-intuitive, but only indicates that the states defined using the different descriptors are not identical although they are correlated.

**Figure 11:**
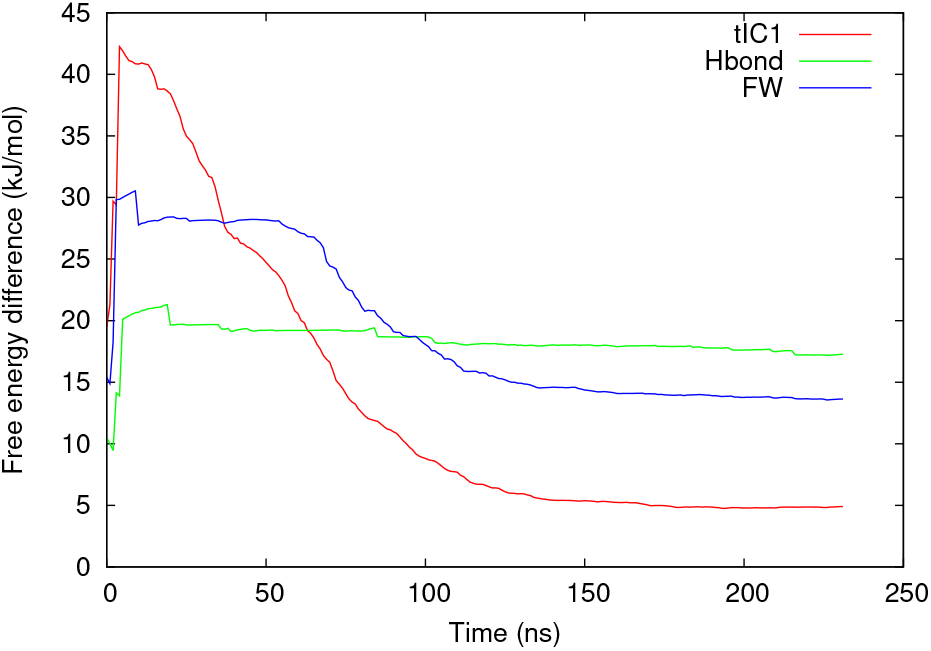
Convergence behavior for the PT-METAD simulation with focus on Ile18. The diagram shows the calculated free-energy difference with respect to tIC1 (right minus left region), Ile18–Tyr35 H-bond distance (broken minus formed), and the Ile18 first-water (FW) distance (solvated minus unsolvated), as a function of simulation time.

The relation between loop motion, hydrogen bonding, and solvent exposure was investigated through two-dimensional free-energy surfaces (Figure 12). It can be seen that the ”right” region (tIC1*>* 0.0028) is more allowing with respect to solvation than the left region, but that the unsolvated substate still dominates. The second diagram shows the metastable state with a longer H-bond distance (∼ 0.37 nm) and a shorter FW distance (∼ 0.22 nm). Its small barrier explains why the broken state is so short-lived for Ile-18 (see e.g. Table 2).

**Figure 12:**
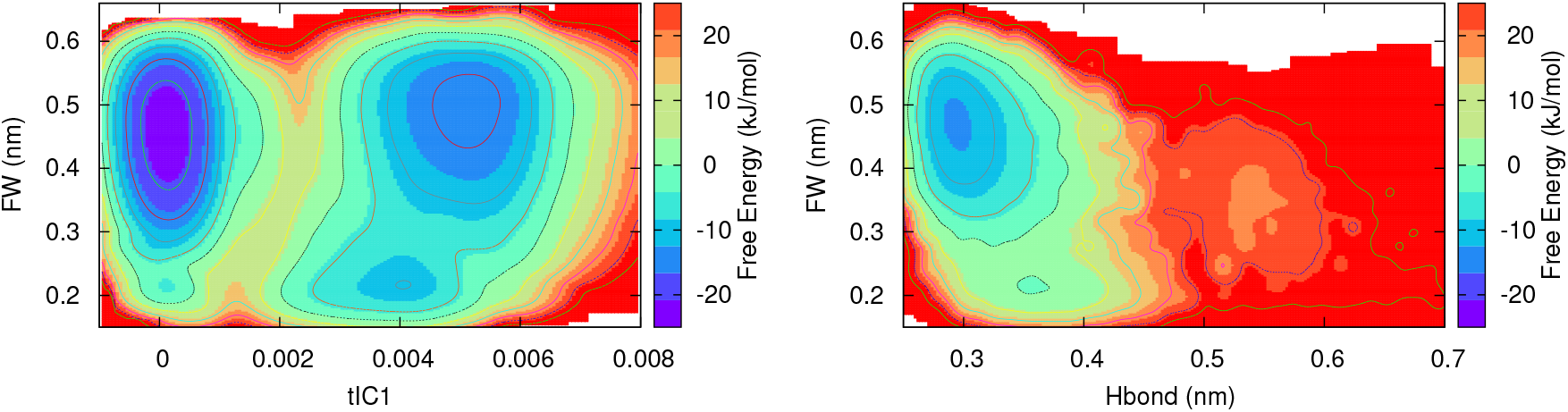
Two-dimensional free-energy surfaces for Ile18 obtained from the PT-METAD simulations. Left: Projection along tIC1 vs. first-water distance (FW). Right: Ile18– Tyr35 N..O distance vs. first-water distance.

### Gly-36

Similar to the Ile18 amide, the DESRES simulation also did a poor sampling of the broken state for Gly36-NH, which is hydrogen-bonded to Thr11-O. The results from PT-METAD is shown in Figure 13. The lack of a clear metastable state could mean that the broken state with its rather long lifetime is not accessed in the PT-METAD simulation but requires movement along some additional slow degree of freedom.

**Figure 13:**
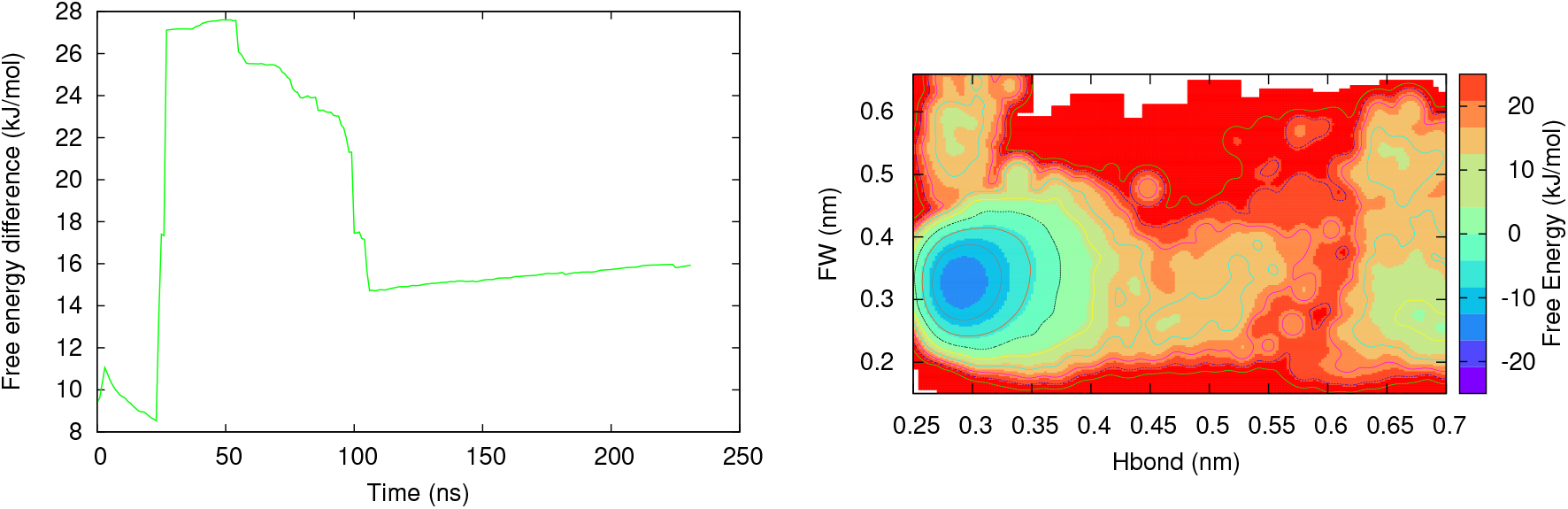
Results for Gly36 extracted from the PT-METAD simulation. Left: Convergence behavior with respect to the free-energy difference along the Gly36–Thr11 hydrogen bond (broken minus formed). Right: Two-dimensional free-energy surface (Gly36–Thr11 N..O distance vs. first-water distance).

#### 3.3.2 Metadynamics with specific CVs

In addition to the PT-METAD simulation with the general tIC1 CV, which mainly enhances the loop motions, we performed simulations with more specific CVs designed to enhance the breaking and reforming of one hydrogen bond at a time. Most of the variants tried during this project used as a CV the H-bond distance transformed by a switching function, in order to focus on the most interesting range (3–4.5 Å) and avoid biasing while inside the broken state. However, after experiencing problems with these switching-function simulations being very sensitive to the parameters, we found that simulations using the distance directly as the CV were in fact more robust, provided that restraining wall potentials were used to limit the exploration of the broken state.

### Ile18

For Ile18, two independent metadynamics simulations were run, starting from the crystal structure and the broken state, respectively. The convergence is shown in Fig. 14 and the two-dimensional FES is shown in Fig. 15. The qualitative features of the 2D FES are the same for the two simulations and involves the same metastable state as was observed in the PT-METAD simulation, as well as another metastable state with completely dissociated hydrogen bond, which is apparently better sampled with the specific CVs. The discrepancy between the quantitative free-energy difference, e.g. with respect to the H-bond distance, predicted by PT-METAD and distance-CV metadynamics (17 and 8–10 kJ/mol, respectively) indicates that further investigation is needed.

**Figure 14:**
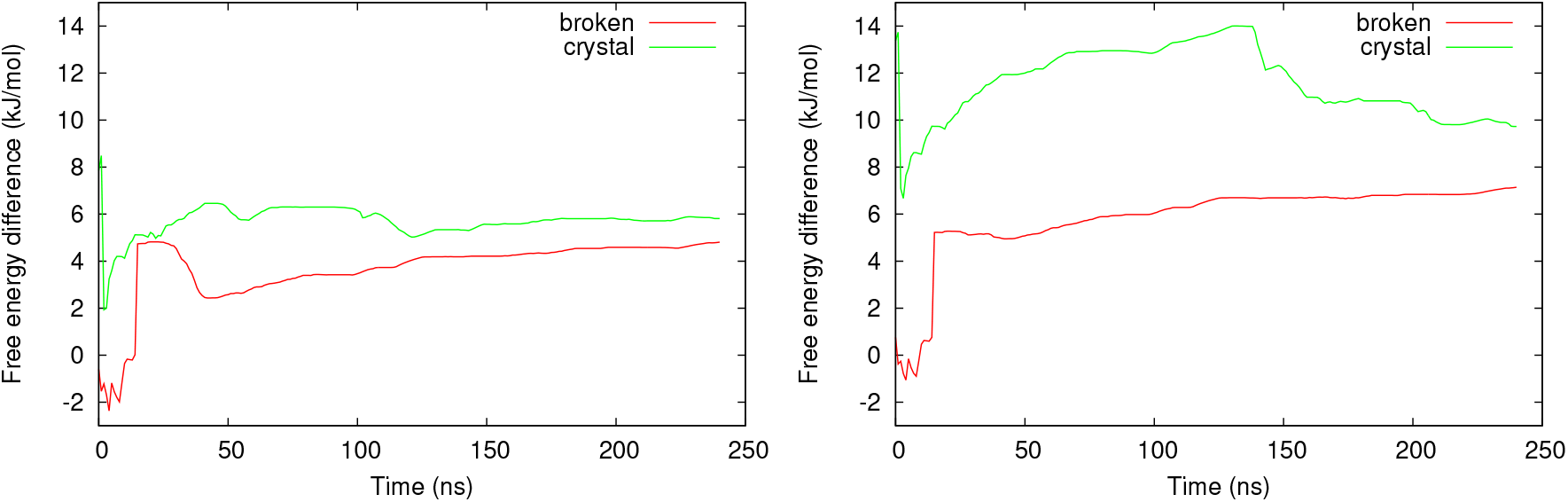
Convergence behavior for distance-CV metadynamics for Ile18. Left: Free-energy difference with respect to H-bond distance (broken minus formed). Right: Free-energy difference with respect to FW distance (solvated minus non-solvated).

**Figure 15:**
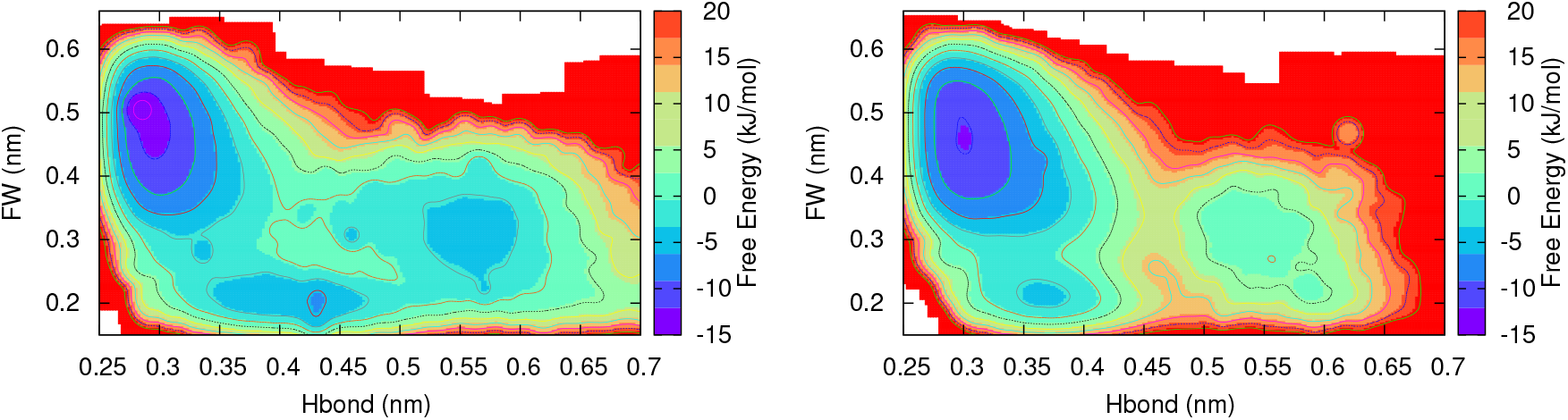
Free-energy surface (H-bond distance versus first-water distance) obtained from distance-CV metadynamics for Ile18. Left: Starting from the broken state. Right: Starting from the crystal structure.

### Gly36

Even for Gly36, two independent metadynamics simulations were run, starting from the crystal structure and the broken state, respectively. The apparent two-dimensional FES for these simulations are shown in Fig. 16, but the lack of agreement clearly illustrates that the two simulations explore different parts of the phase space, probably due to some slow degrees of freedom that were not considered. Nevertheless, both simulations indicate that the free-energy profile along the H-bond distance has more separated basins than in the case of Ile18, so that complete dissociation is needed to reach the metastable state.

**Figure 16:**
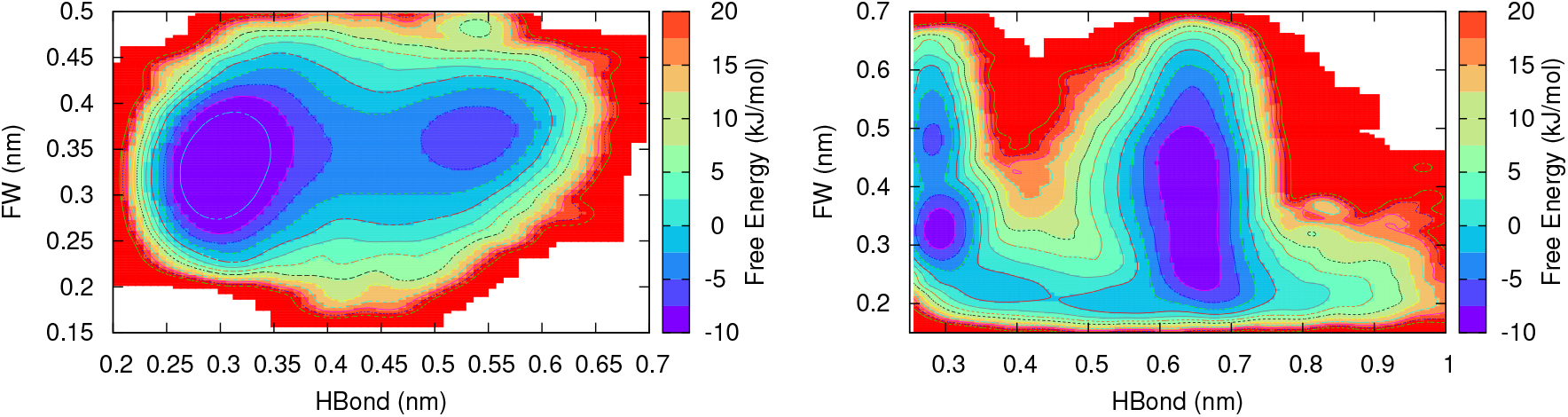
Free-energy surface (H-bond distance versus first-water distance) obtained from distance-CV metadynamics for Gly36. Left: Starting from the broken state. Right: Starting from the crystal structure.

### Met52 and Gly56

The results for Met52 and Gly56 are shown in Fig. 17. Gly56 shows a clear metastable state involving a completely dissociated hydrogen bond and a tightly bound water molecule. The results for Met52 are more inconclusive, but suggest that the broken state is not necessarily solvent-exposed.

**Figure 17:**
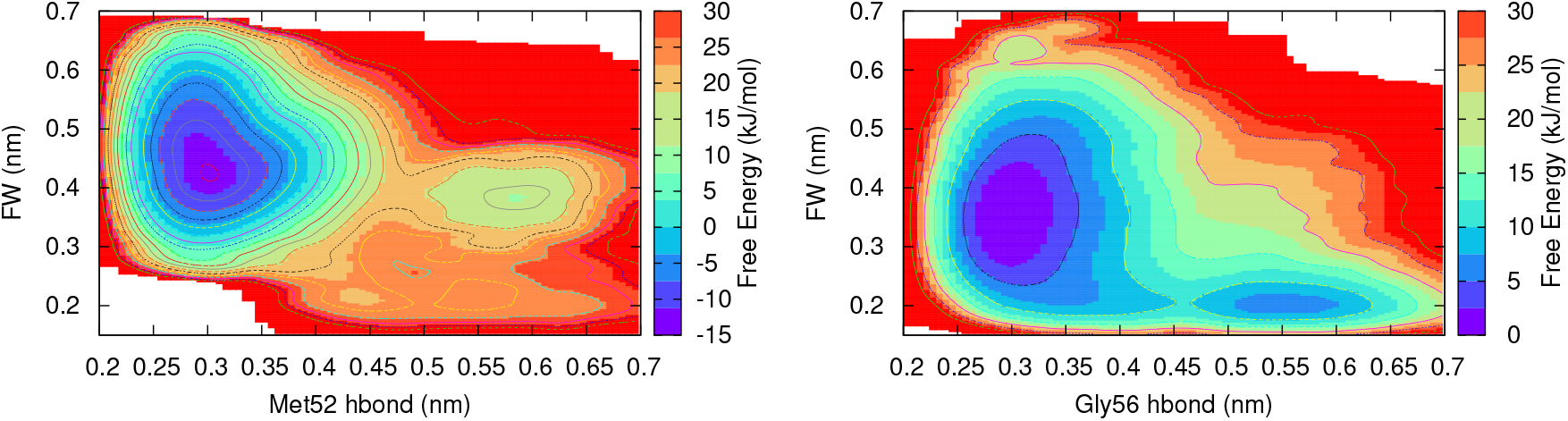
Free-energy surfaces from distance-CV metadynamics for Met52 and Gly56, respectively.

### 3.4 Comparison with experimental results

The reweighted probability distributions from the PT-METAD simulation can be used to estimate the free-energy difference between two arbitrarily defined regions of the conformational space, such as the open and closed states defined in Ref. [4]. Thus, they can also be used to predict protection factors that can be compared to experiment, similarly as was done for a long MD simulation in Ref. [4]. Such comparison is shown in Figure 18.

**Figure 18:**
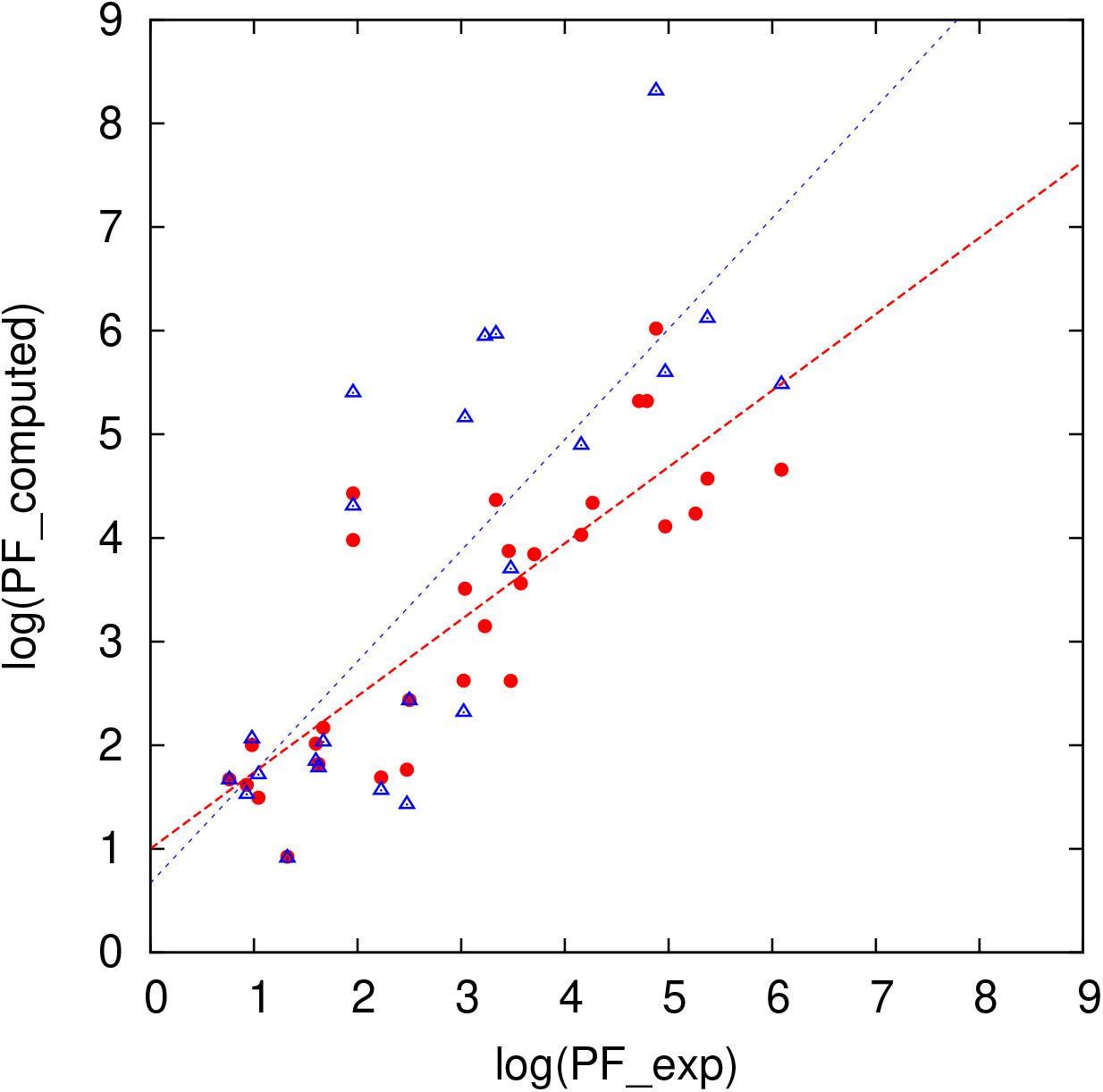
Experimental protection factors (PFs) plotted against PFs estimated from the DESRES trajectory [4] (red circles), and against PFs estimated from the PT-METAD simulation in this study (blue triangles). The corresponding linear fits are also shown.

Given that our enhanced-sampling protocol only assists H-bond breaking for a handful of residues and that our simulation is ∼4000 times shorter, we would expect much worse results than those in Ref. [4]. Therefore, it is interesting to note that the correlation with experiment is not much worse (*R*^2^ = 0.61 instead of 0.68) and the slope of the linear fit is (fortuitously) somewhat better (1.07 instead of 0.74). This is because many amides are so solvent-exposed that a very short simulation suffices to gather statistics, and others are so protected that even the millisecond simulation is too short.

Enhanced sampling with carefully devised CVs should be able to assist the estimation for a few more CVs that exchange via local unfolding. However, many of the amides in the core are protected by a strong network of hydrogen bonds and will only exchange upon complete unfolding of the protein. Such unfolding/folding is not yet accessible by enhanced-sampling approaches for general proteins, although it has been demonstrated for smaller model systems [19].

## 4 Conclusions

The hypothesis that the protein conformation defines the metastable states responsible for hydrogen exchange was confirmed for most residues of BPTI through analysis of the millisecond MD trajectory. For most of the amides with fluctuating solvent exposure, the inflow of one water molecule is almost immediate when the intermolecular hydrogen bond is broken, and the presence of a second water molecule is always a transient state that depends on random fluctuation of the water molecules rather than a distinguishable metastable state.

Enhanced-sampling simulations was used to investigate the nature of the broken states in a more efficient way than classical MD, but it was difficult to find general collective variables that always provide reliable results. For the amide hydrogens that exchange through global unfolding, the problem becomes as difficult as the general folding problem.

## 5 Acknowledgments

We thank D. E. Shaw Research for sharing the BPTI trajectory, Filip Persson for sharing trajectory manipulation scripts, Bertil Halle for inspiring this project, and the Swedish Research Council for financial support.

## Footnotes

1 upper and lower walls are set at 0.8 nm and 0.2 nm respectively

2 upper and lower walls are set at 0.9 nm and 0.2 nm. Respectively

